# The role of the Andean uplift as an asymmetrical barrier to gene flow in the Neotropical leaf-cutting ant *Atta cephalotes*

**DOI:** 10.1101/2021.03.24.436865

**Authors:** Vanessa Muñoz-Valencia, Glever Alexander Vélez-Matínez, James Montoya-Lerma, Fernando Díaz

**Affiliations:** Grupo de Ecología de Agroecosistemas y Hábitats Naturales, Universidad del Valle. Calle 13 No. 100-00, Meléndez 760034, Cali, Valle del Cauca, Colombia; Universidad Nacional de Colombia - Sede Palmira. Carrera. 32 No.12-00, Chapinero 763536, Palmira, Valle del Cauca, Colombia; Department of Entomology, University of Arizona, Tucson, AZ, USA

**Keywords:** Amazon, Andes mountains, Colombia, gene flow, isolation by distance, isolation by barrier, leaf-cutting ants, Neotropics

## Abstract

Neotropical diversification by the Andean uplift is typically addressed on a large evolutionary scale (*e*.*g*. speciation), even though many species are still distributed in both sides of the mountains. The three parallel mountain ranges in the northern Andes (Colombia) impose a major geographical barrier to species’ migration from South to Central America. How important these barriers are for conspecific diversification of cross-Andean species such as the leaf-cutting ants remains largely unknown. To answer this question, we studied the *mtCOI* gene of *Atta cephalotes*, the most widely distributed leaf-cutting ant species. Our hierarchical analyzes evidenced substantial genetic structure among regions and populations, suggesting a more complex biogeographical history of Andean populations than previously thought. These mountains seem to isolate Central American and Western Colombian populations from the rest of *A. cephalotes* in South America. Population and migration modelling are consistent with the origin of this species in South America and a major role of the Eastern cordillera as a geographical barrier to historical gene flow, restricting dispersion from north to south. These findings provide insights into the role of the Andean uplift as barrier to gene flow and, eventually, implications for monitoring and designing management strategies for leaf-cutting ants.

## 1 INTRODUCTION

There is a wealth of evidence demonstrating that the Andean uplift has promoted allopatric diversification in a wide range of taxa (Hoorn et al., 2010; Pérez-Escobar et al., 2017; Salgado-Roa et al., 2018). These mountains reach their maximum complexity in northern South America (Colombia), forming a branch of three parallel mountain ranges separating flora and fauna in a North-South manner (Gustavo H. Kattan, Franco, Rojas, & Morales, 2004; Luebert & Weigend, 2014). This pattern of diversification is common to mammals (Antonelli et al., 2009; Hoorn et al., 2010), birds (Antonelli et al., 2009; Brumfield & Capparella, 1996; Cadena, Pedraza, & Brumfield, 2016), arthropods (Hoorn et al., 2010; Salgado-Roa et al., 2018), and plants (Lagomarsino, Condamine, Antonelli, Mulch, & Davis, 2016; Luebert & Weigend, 2014; Pérez-Escobar et al., 2017). Several hypotheses have been discussed for decades to explain neotropical biodiversity distributions, often calling for Pleistocene refugia (Antonelli et al., 2009; Solomon, Bacci, Martins, Vinha, & Mueller, 2008) or vicariance mechanisms (Harrison, 1991; Miller et al., 2008; Salgado-Roa et al., 2018) as alternative hypothesis linked to the Andean uplift. More recently, genetic data suggested that diversification occurred through a range of more complex mechanisms across the timeline of the uplift history than initially thought (Cadena et al., 2016; Pérez-Escobar et al., 2017).

The impact of the Andean uplift on neotropical evolution seems to rely on the life history and species’ ecological context, with birds generally showing more recent genetic divergences when compared with other vertebrates (Cadena et al., 2016; Hoorn et al., 2010). The divergence spectrum across the Andes mountain ranges from restrictions to conspecific gene flow in some species (Masello et al., 2011; Milá, Wayne, Fitze, & Smith, 2009), to multiple independent vicariant events (Chapman, 1917). Although the evidence for insects is more limited, a few studies in butterflies from Northern Andes (De-Silva et al., 2017; Elias et al., 2009) and bees and stick-insects in Southern Andes (Dick, Roubik, Gruber, & Bermingham, 2004) support multiple isolating mechanisms rather than a single geoclimatic event. Much of this evidence comes from comparative species phylogenies with more ancient implications (*e*.*g*. speciation). The role of these mountains in conspecific diversification remains more elusive, even though many species still distribute in both sides of their putative physical barriers (Dick et al., 2004; Salgado-Roa et al., 2018) suggesting recent diversification or gene flow still connecting conspecific populations. These mechanisms are better detected in species with a wide distribution range across the Andes (Cadena et al., 2016; Dick et al., 2004), such as the leaf-cutting ants, whose entire biogeographical history has occurred across this Neotropical landscape.

Previous studies investigating the origin and dispersion by leaf-cutting ants (Mayhé-Nunes & Jaffé, 1998; Mueller et al., 2017; Solomon et al., 2008; Wetterer, Schultz, & Meier, 1998), have been focusing on species with limited distributions or are based only on part of their ranges. Although this topic remains unresolved, the most convincing hypothesis place the origin of the leaf-cutting ants in subtropical areas (grasslands) of South America, gradually extended across the continent and later spread into Central America (Chapela, Rehner, Schultz, & Mueller, 1994; Mueller et al., 2017; Solomon et al., 2008; Weber, 1972). A recent study has challenged this initial hypothesis proposing a Central American origin for the evolution of *Atta* agriculture, with secondary dispersion into South America (Branstetter et al., 2017). However, there is no fossil evidence supporting the presence of the group in Central America before the formation of the Panama land bridge (Mueller et al., 2017). Heríe we explored both hypotheses by considering the evolutionary history of *Atta cephalotes*, the most widely distributed Attini species in the Neotropics (Della Lucia, Gandra, & Guedes, 2014; Fernández, Castro-Huertas, & Serna, 2015) to investigate patterns of gene flow across the northern Andes. This species ranges from southern Mexico to northern Argentina, and from the Lesser Antilles to the north of Barbados (Fernández et al., 2015; Fernández & Sendoya, 2004). Although *A. cephalotes* is common in both natural and rural habitats, this species has expanded to novel environments as a response to multiple ecological and anthropic factors, causing important economic consequences when nesting near cropping and forest systems or urban constructions (Chacón de Ulloa, 1994; Leal, Wirth, & Tabarelli, 2014; Montoya-Lerma, Giraldo-Echeverri, Armbrecht, Farji-Brener, & Calle, 2012).

To our knowledge, Solomon et al. (2008) reported the solely study comparing genetic data from *Atta* species on a large scale in the Neotropics. Only a few *A. cephalotes* samples from northern Andes were included (although not representative samples). Interestingly, such samples showed a closer relationship to Central rather than to southern South American samples. The three Andean cordilleras are located in this South-to-North transitional land in Colombia, suggesting their role in this species’ genetic divergence. This indicates that insect populations around these mountains might reveal genetic evidence to elucidate the role of the Andean uplift in the evolutionary history of these ants as well as provide evidence in support of any of the two origin hypotheses for this species. We investigate these questions in *A. cephalotes* using *mtCOI* gene sequences sampled in several regions across the three Andean cordilleras in Colombia. We demonstrate that the Eastern cordillera represents a geographical barrier to gene flow, restricting migration from Central America and Western Colombian populations to the rest of South America.

## 2. METHODS

### 2.1 Sampling

We sampled *A. cephalotes* workers from nests in four different Colombian regions: The Pacific region, the Andean region (sub-divided into two smaller climatically distinct sub-regions: Andean_1 between 800 - 1,050 m.a.s.l. and Andean_2 between 1,300 - 2,200 m.a.s.l.), and the Amazonian region, for a total of 164 sampled ants. These are the main climatic regions in the transition lands between South and Central America of *A. cephalotes’* distribution, which were analyzed together with 42 additional *mtCOI* sequences from different locations in Central and South America obtained from GenBank (Figure 1), for a total of 206 sequences in 23 locations between South and Central America. We considered one sample per nest that were at least 1.5 km apart from each other in order to reduce the probability of sampling non-independent nests. Species were identified using taxonomic keys following Fernández et al. (2015). Vouchers were kept at the Entomological Museum at the Universidad del Valle (MUSENUV), Cali, Colombia.

**Figure 1.**
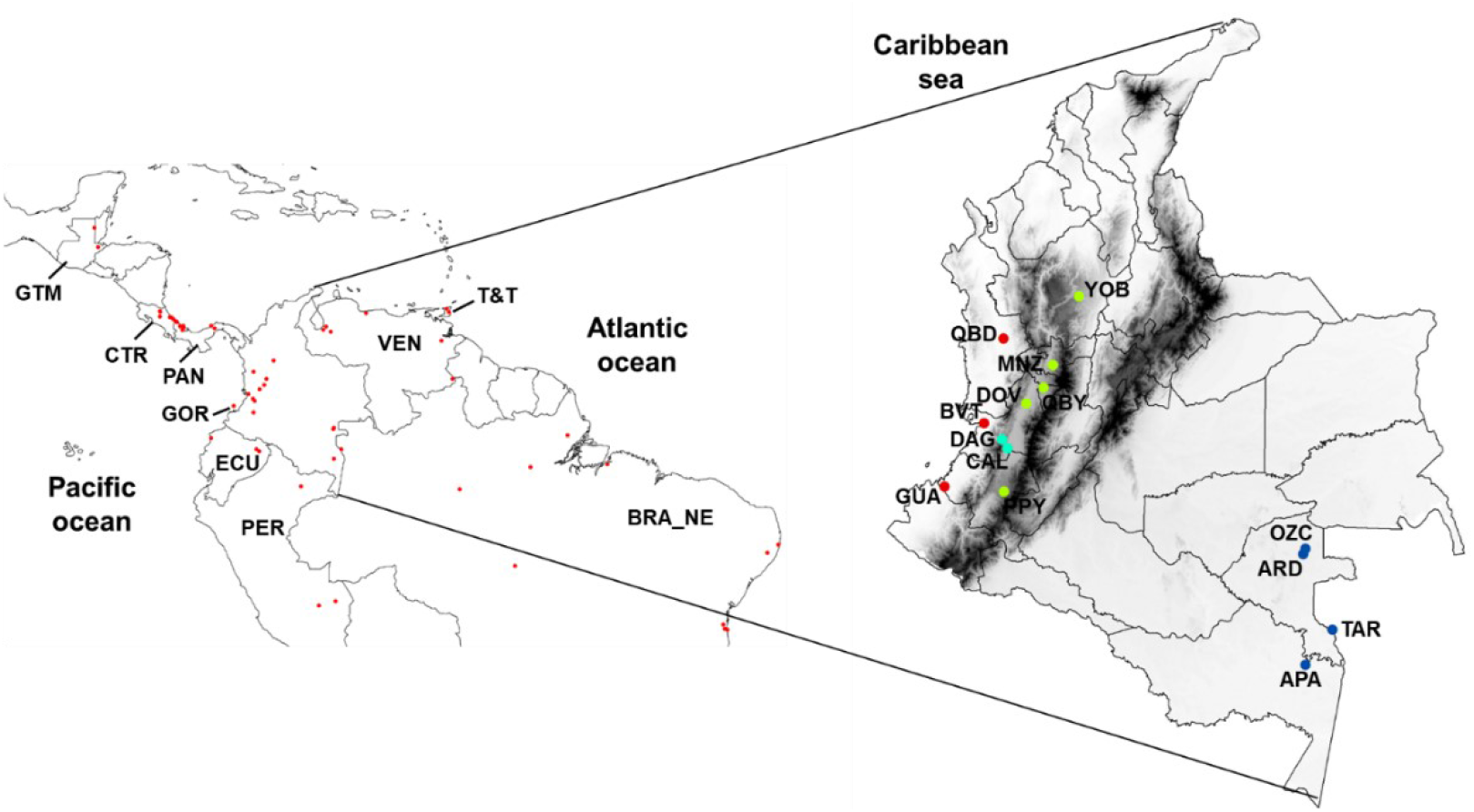
Study area, including 24 sampling localities in Neotropical regions of *A. cephalotes*’ distribution. BVT, Buenaventura; QBD, Quibdó; DAG, Dagua; GUA, Guapi; PPY, Popayán; GOR, Gorgona; DOV, El Dovio; CAL, Cali; QBY, Quimbaya; YOB, Yolombó; MNZ, Manizales; APA, Apaporis; ARD, Arrendajo; OZC, Ozcimi; TAR, Taraira; CTR, Costa Rica; GTM, Guatemala; PAN, Panama; BRA, Brazil; BRA_NE, northeastern Brazil; T&T, Trinidad and Tobago; ECU, Ecuador; PER, Peru; VEN, Venezuela.

### 2.2 DNA extraction

Ants were numbed with cold, washed in 70% ethanol for two minutes to remove possible external contaminants and then stored in 96% ethanol at -20°C until DNA extraction. Total DNA was extracted from individual samples using TNES lysis buffer pH 7.5 (Tris 50 mM, NaCl 0.4 M, EDTA 100 mM, SDS 0.5%) and chloroform: isoamyl alcohol (24:1). Whole ants were homogenized in 50 µl of TNES buffer, followed by addition of 500 µl of TNES buffer and 10 µl of proteinase K (20 mg/ml). Samples were incubated over-night at 56°C and then proteinase K was inactivated at -20°C for 15 min. RNA was depleted using 5 µl of RNase A (10 mg/ml) at 37°C for 30 min followed by RNase A inactivation at -20°C for 15 min. The protein solution was then precipitated using 200 µl of NaCl 5 M and then centrifuged at 15,000 g for 7 min. The supernatant was transferred to a new tube containing 600 µl of chloroform: isoamyl alcohol solution (24:1) and centrifuged at 15,000 g for 6 min. The supernatant was finally transferred to a new tube and DNA was pelleted with 600 µl of isopropanol at 15,000 g for 5 min and then washed using 600 µl of 70% cold ethanol followed by centrifugation at 15,000 g for 5 min. The supernatant was discarded, and the DNA pellet was dried at 45°C for 15 min, eluted with ultrapure water and stored at -20°C. DNA quality and quantity were estimated using a NanoDrop 2000 spectrophotometer (Thermo Scientific).

### 2.3 PCR and sequencing

A *mtCOI* fragment of approximately 380 bp was amplified, using universal primers for the *mtCOI* gene (Ben – 5’ GCTACTACATAATAKGTATCATG 3’ and Jerry – 5’ CAACATTTATTTTGATTTTTTGG 3’)

as suggested by Kronauer et al. (2004) and Simon et al. (1994). PCR reactions were performed in 10 µl volume, consisting of 2 µl of DNA, GoTaq® Master Mix (PROMEGA) 1X and 0.2 µM of each primer, using the following cycling conditions: 94°C for 2 min, 30 cycles of 94°C for 1 min, 58°C for 1, and 72°C for 1 min, followed by a final extension at 72°C for 10 min. PCR products were confirmed by 1% agarose gels and sent to Macrogen Corp. USA for sequencing. The quality of electropherograms was analyzed with Geneious 7.1.3 (https://www.geneious.com), excluding sequences with low-quality and ambiguous bases. Sequences were confirmed as belonging to *A. cephalotes* using BLASTn (NCBI, available online). Obtained sequences were aligned using the *CLUSTAL W* algorithm in BioEdit running the default parameters (Tamura, Stecher, Peterson, Filipski, & Kumar, 2013). Aligned sequences included 42 from Central and South America regions that were obtained from GenBank as well as outgroup taxa: *A. colombica, A. laevigata, A. sexdens* and *Acromyrmex striatus* (Figure 1). All newly generated sequences of *A. cephalotes* were deposited in GenBank (GenBank accession numbers: MW245066 - MW245234).

### 2.4 Genetic diversity and demographic history

Only locations with sample size n ≥ 5 nests were included in this analysis (191 samples in 17 locations) to reduce sampling bias while estimating population structure. Genetic diversity parameters were estimated for locations and regions in DnaSP v.5.19 (Librado & Rozas, 2009) and were summarized as the number of polymorphic sites (S), the number of haplotypes (h), nucleotide diversity (π) and haplotype diversity (h_d_). Fu’s Fs (Fu, 1997) and Tajima’s D (Tajima, 1989) neutrality tests were used to assess evidence of recent population expansion when the null hypothesis of neutrality was rejected due to significant negative values (*P* < 0.02 for Fs and *P* < 0.05 for D). Further, Ramos-Onsins & Rozas R^2^ and raggedness (r) neutrality tests were computed to detect population growth (Ramos-Onsins & Rozas, 2002). Fu’s FS performs better for large sample sizes whereas R^2^ test is superior for small sample sizes (Ramos-Onsins & Rozas, 2002). Demographic history was also examined by analyzing distribution of pairwise differences between individual sequences through mismatch distribution in DnaSP v.5.19. A haplotype network was constructed using a median-joining algorithm in Network 5.0.0.3 (Bandelt, Forster, & Rohl, 1999) for the whole data set (set of 206 samples).

### 2.5 Population structure

As described for genetic diversity analysis, only locations with sample size n ≥ 5 were included in this analysis (191 samples in 17 locations). A hierarchical analysis of molecular variance (AMOVA) was performed in ARLEQUIN v.3.5.2.2 (Excoffier & Lischer, 2010) to evaluate population structure at two hierarchical levels (regions and populations within regions), as well as estimating pairwise fixation indices (*Φ*_ST_) between populations. Further, pairwise genetic distances were calculated using MEGA X based on the Tamura 3-parameter model. This matrix was used to examine the pattern structure among samples using the UPGMA algorithm (unweighted pair group method using an arithmetic average) in MEGA X. A discriminant analysis of principal components (DAPC) was also performed on individual haplotype frequencies using the R package ADEGENET (Jombart, 2008; Jombart, Devillard, & Balloux, 2010) to represent clusters of genetically related individuals (Jombart et al., 2010). This analysis requires data transformation using a principal component analysis (PCA) as a previous step to a discriminant analysis (DA). For DA, the “*dapc*” function was implemented in order to determine the optimal number of PCs for the DAPC based on their cumulative variance. The “*scatter*” function was used to plot the results as a scatter plot. Pairwise genetic distances were also used to assess the divergence across *mtCOI* sequences. Divergence times among regions were estimated using a conventional *mtCOI* clock of 3% sequence divergence between a pair of lineages per million years (Hebert, Cywinska, Ball, & Waard, 2003). Here, we grouped the whole set of 206 samples by regions as follow: Western Colombia (Pacific, Andean_1, and Andean_2), Amazonian, northeastern Brazil (Brazil_NE) and Central America regions and then pairwise genetic distances and divergence times were calculated.

A Spatial Analysis of Molecular Variance (SAMOVA) was performed using SAMOVA v.1.0 (Dupanloup, Schneider, & Excoffier, 2002) in order to identify major spatial clusters in analyzed regions by accounting for their geographical distribution while analyzing their genetic differentiation (Dupanloup et al., 2002). For this, populations groups that are phylogeographically homogeneous and maximally differentiated by geographic distance were identify. We estimated the maximum plateau value of *F*CT through the space distribution of compared populations in order to select the optimal number of groups (K) (Shi, Kerdelhué, & Ye, 2012). This analysis was performed for K values from 1 to 8 in order to identify the most likely number of groups amongst the 17 populations characterized by *mtCOI*.

### 2.6 Scenarios of genetic differentiation – IBD vs IBB

We exploited the power of Redundancy Analysis (RDA) in order to disentangle the relative explanatory contributions of alternative scenarios of genetic divergence across the *A. cephalotes*’ range. RDA has been highly suggested as a powerful method for reflecting a linear spatial relationship of *Φ* _ST_ and gene flow [as opposed to traditional Mantel test (Bradburd, Ralph, & Coop, 2013; Legendre & Fortin, 2010; Meirmans, 2015)]. We performed a distance-based RDA (db-RDA) by computing the influence of spatial (geographic distance matrix) and topography (barrier) variables on pairwise genetic distance matrix (*Φ*_ST_). This is performed by combining Principal Component Analysis (PCA) and multiple regression using Euclidean distances (Meirmans, 2015; Wang, Glor, & Losos, 2013).

An RDA-based model comparison was implemented considering alternative scenarios of genetic isolation. The model with the Andes as a barrier to gene flow, isolation by barrier (IBB), was compared against a model of isolation by distance (IBD). Explanatory variables were grouped into two classes according to their resulting pattern of isolation. Variables representing geographical distance between populations (space) for IBD; and those representing the Andes mountains as a geographical barrier splitting populations in an allopatric manner for the IBB model. RDA models were then used to identify the best ordination model that describes genetic differentiation (James, Coltman, Murray, Hamelin, & Sperling, 2011).

To test for isolation by distance (IBD), pairwise geographical distances expressed as a distance matrix were transformed into a vector format (De Queiroz, Torrente-Vilara, Quilodran, da Costa Doria, & Montoya-Burgos, 2017; Noguerales, Cordero, & Ortego, 2016) using the ‘*pcnm’* function in the R package VEGAN v.2.2.0 (Oksanen et al., 2019). Then, spatial explanatory components were tested through a db-RDA analysis and only the significant explanatory PCNM components were retained. This analysis was conducted using the “*ordistep”* function of the VEGAN package, to prevent over fitting. Selected components were used to test for IBD through a final db-RDA against pairwise *Φ*_ST_ using the “*capscale*” function of the same package. IBB was tested using an *Andes classification* of populations represented by a dummy variable using two different classifications. First, populations were split up into three sections following the Colombian Andes cordilleras as independent barriers: (0) for populations located on the west side of the Western Andes range, (1) for populations located in the east side of this range, and (2) for populations located in the east side of the Eastern cordillera. Second, populations were split up into two sections according to the barrier imposed only by the Eastern cordillera: (0) for populations from Central America and Western Colombia (Pacific and Andean regions) located on the west side of this range, and (1) for populations located in the east side, the rest of South America (Figure 1). Significance of predictors was assessed using multivariate *F*-statistics with 9999 permutations using the ‘*anova*.*cca*’ function included in the package VEGAN. All explanatory variables were scaled using the ‘*scale’* function in the package VEGAN. The whole model including both IBD and IBB predictor variables was compared to the null model (*i*.*e*. intercept), to define the best combination of variables to be included into the model selection according to the AIC criterion. Model selection was performed using the “*ordistep”* function of the package VEGAN, to prevent over fitting. Once all models of genetic isolation were defined and evaluated, the best model involving IBB was tested through a conditional partial RDA (conditional test), accounting for IBD. Finally, the relative contribution of these models to explain patterns of gene flow in *A. cephalotes* was evaluated in the final model, including all mechanisms of genetic isolation using a variation partitioning analysis with the function ‘*varpart*’ implemented in the R package VEGAN. This analysis allowed us to quantify the relative contributions of IBB vs IBD in shaping the distribution of genetic diversity across *A. cephalotes*’ distribution.

### 2.7 Migration estimates

MIGRATE v4.x software (Beerli, Mashayekhi, Sadeghi, Khodaei, & Shaw, 2019) was used to investigate the direction of gene flow between major genetic groups across the Andes: Western Colombia –WC, Central America –CA and Amazonian-South America –AS by using MCMC simulations. Twenty replicates were used per simulation with a burn-in of 100,000 steps followed by 100 million steps sampled every 100th. Each replicate was run under a static heating approach implementing four incrementally heated chains. Analyses were performed in two sets of model selection following migration patterns based on their Marginal likelihoods, Bayes factors and posterior probabilities (Beerli et al., 2019). Given that WC and CA are more closely related when compared to AS, we first used MIGRATE to confirm whether this is the most likely topology in a set of 9 population models in full Isolation-with-migration. This result was then used in a second set of 15 simulations involving different combinations in order to investigate the direction of gene flow across the Andes mountains in *A. cephalotes*.

## 3. RESULTS

### 3.1 Haplotype network

Thirty-seven *mt*COI haplotypes were found in the whole data set of 206 sequences, many of which were population- or region-specific. Western Colombian regions shared a few haplotypes with Central American region (Hap1, Hap2, Hap4, and Hap7). The Amazonian region shared only one haplotype with Central America and northeastern Brazil (Hap11). Twenty of the population-specific haplotypes were exclusively found in one individual per population. The number of nucleotide differences between haplotypes sequences detected ranged from 1 to 17 bp. Two haplotypes from Brazil were the most divergent, with 17 and 13 mutations, respectively, suggesting that southern South America ant populations not only contain the highest genetic diversification within *A. cephalotes* but are also the most divergent.

The haplotype network suggests three different genetic groups (I-III). The first group (I) corresponds to samples from the Amazonian region (except for haplotype Hap5). The second group (II), with two main haplotypes Hap1 and Hap4, included most samples from Western Colombia – WC and Central America – CA regions (Figure 2). In this group, Hap1 was the most common haplotype, represented by 38% of the samples, which were dominant in WC and CA. Hap4 was detected in 10% of the samples, being exclusive to Western Colombia (Figure 2). The third group (III) was composed by northeastern Brazilian sequences.

**Figure 2.**
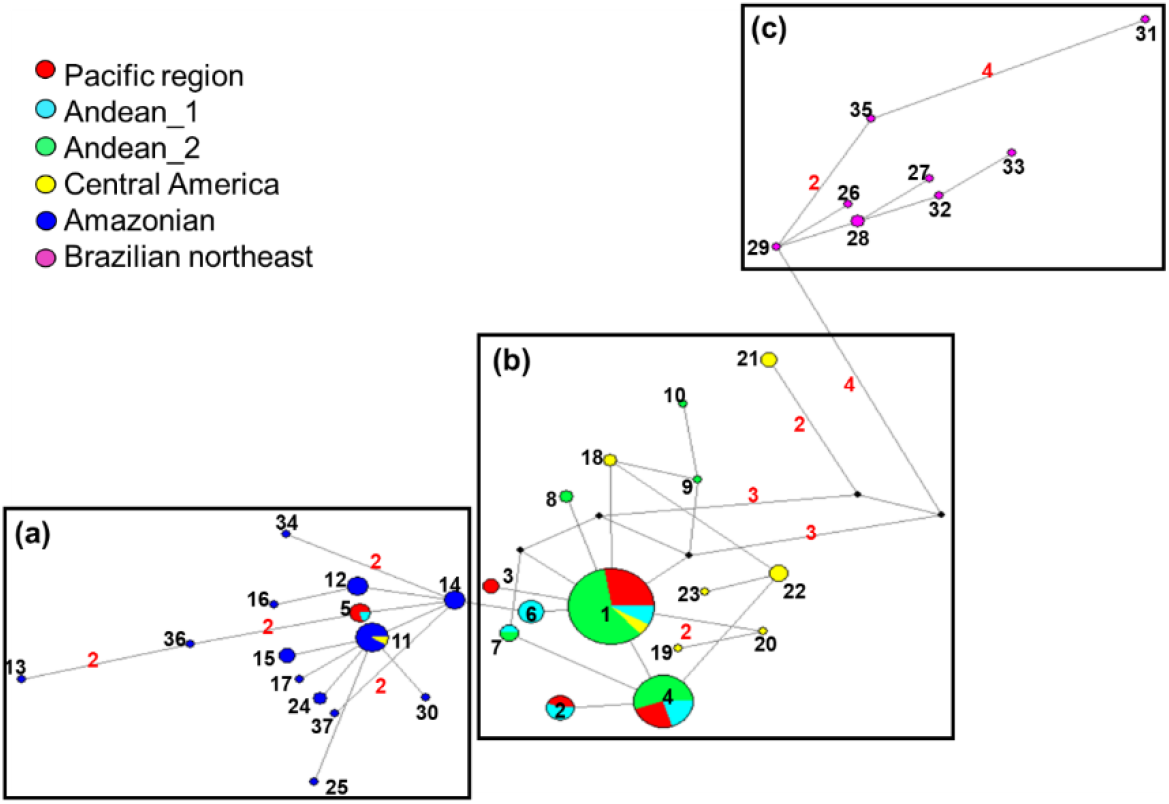
Haplotype network based on sequences of the *A. cephalotes mtCOI* gene sampled in six different Neotropical regions from South to Central America. Median-joining network based on 206 samples. Circles represent each haplotype, and the size of each circle is proportional to the number of individuals within each haplotype. Red numbers connecting lines indicate the number of mutations; lines without number indicate one mutation. Black circles represent missing intermediate haplotypes. Boxes represent haplotype groups per region: (a) corresponds with Amazonian haplotypes except for Hap5; (b) included Western Colombia and Central America haplotypes; (c) was represented by Northeastern Brazil haplotypes. Haplotypes are shaded according to region representation.

#### Genetic diversity and demographic history

The final dataset included 191 sequences in 17 populations after filtering for minimum sample size (n ≥ 5). Thirty-five variable sites with 36 mutations were found defining a total of 32 haplotypes in this set. Global nucleotide diversity (π) was 0.9%, while within-population π ranged from 0.0 to 1.1% (Table 1). Haplotype diversity (Hd) in the entire data set was 78%, and within population ranged from 0.0 to 100%. Amazonian, Central America, and northeastern Brazil regions showed the highest diversity for both π and Hd (Table 1) when compared to Western Colombian regions (Pacific, Andean_1, and Andean_2 regions). No evidence for population expansion at regional and population scales were found according to Tajima’s D and Fu’s FS values (Table 1). Moreover, these results were supported by mismatch distributions, showing a multimodal distribution for both overall and location levels, as well as Ramos-Onsins & Rozas (R2) and raggedness (r) neutrality tests.

**Table 1.** Molecular diversity parameters and neutrality tests of the *mtCOI* gene in *Atta cephalotes* populations form South to Central America.

| Region/Location | n | S | h | % H<sub>d</sub> | k | % $\pi$ | D | P<sub>D</sub> | FS | P<sub>FS</sub> |
| --- | --- | --- | --- | --- | --- | --- | --- | --- | --- | --- |
| **Pacific** |  |  |  |  |  |  |  |  |  |  |
| BVT | 9 | 2 | 2 | 22 | 0.44 | 0.2 | -1.36 | 0.08 | 0.67 | 0.48 |
| QBD | 13 | 5 | 4 | 60 | 1.36 | 0.4 | -0.56 | 0.32 | 0.16 | 0.51 |
| GUA | 20 | 5 | 4 | 56 | 1.33 | 0.4 | -0.17 | 0.45 | 0.68 | 0.65 |
| Region total | 42 | 6 | 5 | 67 | 1.24 | 0.4 | -0.29 | 0.41 | 0.31 | 0.60 |
| **Andean_1** |  |  |  |  |  |  |  |  |  |  |
| DAG | 10 | 4 | 3 | 51 | 1.11 | 0.4 | -0.82 | 0.25 | 0.68 | 0.62 |
| CAL | 19 | 3 | 4 | 76 | 1.06 | 0.4 | 0.65 | 0.78 | 0.08 | 0.52 |
| Region total | 29 | 6 | 6 | 82 | 1.50 | 0.5 | -0.06 | 0.52 | -0.51 | 0.38 |
| **Andean_2** |  |  |  |  |  |  |  |  |  |  |
| PPY | 14 | 2 | 3 | 62 | 0.70 | 0.3 | 0.32 | 0.70 | 0.15 | 0.42 |
| DOV | 19 | 2 | 3 | 29 | 0.30 | 0.1 | -1.12 | 0.20 | -1.15 | 0.06 |
| QBY | 15 | 1 | 2 | 25 | 0.25 | 0.1 | -0.40 | 0.30 | 0.13 | 0.29 |
| YOB | 16 | 0 | 1 | 0 | 0.00 | 0.0 | 0.00 | 1.00 | 0.00 | N/A |
| MNZ | 11 | 3 | 3 | 35 | 0.84 | 0.3 | -0.63 | 0.29 | 0.24 | 0.45 |
| Region total | 75 | 6 | 6 | 53 | 0.68 | 0.2 | -0.48 | 0.39 | -1.83 | 0.19 |
| **Amazonian** |  |  |  |  |  |  |  |  |  |  |
| ARD | 5 | 6 | 3 | 80 | 2.60 | 0.9 | -0.67 | 0.35 | 1.09 | 0.70 |
| OZC | 5 | 4 | 4 | 90 | 2.20 | 0.7 | 0.96 | 0.81 | -0.85 | 0.16 |
| TAR | 5 | 4 | 5 | 100 | 1.80 | 0.6 | -0.41 | 0.45 | -3.30 | 0.03 |
| VNZ | 7 | 6 | 3 | 52 | 1.91 | 0.6 | -1.12 | 0.17 | 1.22 | 0.78 |
| Region total | 22 | 11 | 9 | 87 | 2.18 | 0.7 | -0.96 | 0.19 | -2.65 | 0.06 |
| **Central America** |  |  |  |  |  |  |  |  |  |  |
| PAN | 6 | 4 | 3 | 73 | 1.53 | 0.5 | -0.68 | 0.34 | 0.54 | 0.54 |
| CTR | 9 | 12 | 5 | 81 | 4.50 | 1.5 | 0.09 | 0.56 | 0.96 | 0.70 |
| Region total | 15 | 13 | 7 | 87 | 3.51 | 1.1 | -0.49 | 0.32 | -0.26 | 0.45 |
| **Southern South America** |  |  |  |  |  |  |  |  |  |  |
| BRA_NE | 8 | 8 | 7 | 96 | 2.96 | 1.0 | -0.19 | 0.43 | -3.28 | 0.02 |
| Region total | 8 | 8 | 7 | 96 | 2.96 | 1.0 | -0.19 | 0.43 | -3.28 | 0.02 |

Sample size (n); number of variable sites (S); number of haplotypes (h); haplotype (H<sub>d</sub>) and nucleotide ( $\pi$ ) diversities; average number of nucleotide differences (k); Neutrality test: Tajima's D (D) and Fu's FS (FS) and corresponding significance level at $\alpha = 0.05$ (P<sub>D</sub> and P<sub>FS</sub>) after Bonferroni correction. Cannot be calculated (N/A): only one allele in the sample.

### 3.2 Genetic structure of populations

The AMOVA revealed strong population differentiation at all hierarchical levels (Table 2). Most of the genetic differentiation was due to regional effect (∼49%), followed by population sub-structure (∼12%). Overall *Φ*_ST_ was high (*Φ* _ST_ = 0.61) and pairwise *Φ*_ST_ ranged from 0.00 to 0.97, with significant genetic differentiation amongst 115 out of 136 comparisons (Figure 3A). Such comparisons evidenced that regional effect is strongest between the Amazonian and northeastern Brazil regions relative to Western Colombia and Central America (Figure 3A). This is consistent with the Median-Joining network results.

**Table 2.** AMOVA for *Atta cephalotes* populations from South to Central America. Results are shown following our regional classifications as well that suggested by SAMOVA results (Figure 3B).

| Source of variation | Without pops $n \leq 5$ | | | Based on SAMOVA | | |
| --- | --- | --- | --- | --- | --- | --- |
|  | df | Fixation indexes | % of variation | df | Fixation indexes | % of variation |
| Regions | 5 | $\Phi_{CT}$: 0.49 | 49* | 2 | $\Phi_{CT}$: 0.73 | 73* |
| Populations (Regions) | 11 | $\Phi_{SC}$: 0.22 | 12* | 14 | $\Phi_{SC}$: 0.23 | 6* |
| Within pops | 174 |  | 39* | 174 |  | 21* |
| **Total** | 190 |  | 100 | 190 |  | 100 |

\*Indicates significant values, $P \leq 0.001$ .

**Figure 3.**
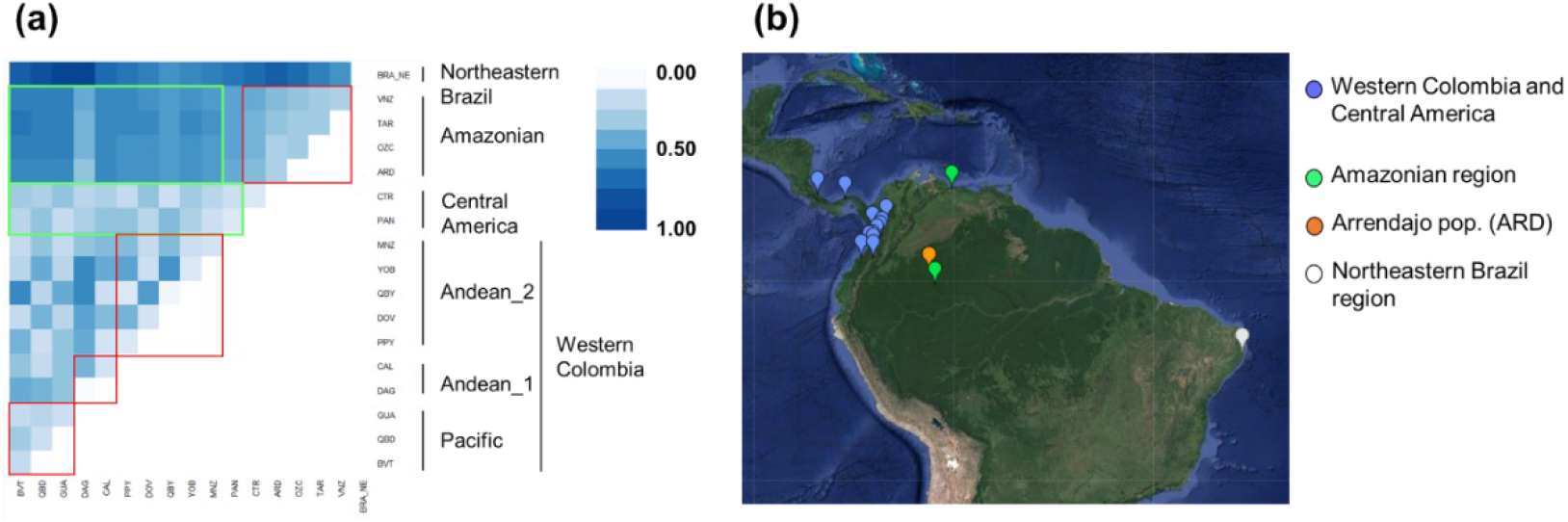
Genetic differentiation of *A. cephalotes* populations. (a) Heat map showing pairwise *Φ* _ST_ values based on *mtDNA* data of 17 populations of *A. cephalotes*. Red squares indicate intra-region comparisons, while at inter-region comparisons between Amazonian and Central America vs Western Colombia are represented by green squares between. The heat map color indicates the level of genetic differentiation, Central American and Western Colombian populations are more closely related when compared with the rest of southern South America (northeastern Brazil and Amazonian). (b) SAMOVA clustering for *A. cephalotes* populations. Color codes of populations correspond to the three groups (one single population, ARD) identified by SAMOVA inferred from 191 mitochondrial sequences (17 *A. cephalotes* populations).

The spatial analysis of molecular variation (SAMOVA) identified three spatial clusters (K = 3), including one single-population (ARD; Figure 3B). These three main groups were geographically consistent and correspond to the same clusters detected by the haplotype network, population structure, and clustering methods (Figures 2 and 4). A second AMOVA was further performed following the observed clustering detected by SAMOVA (while ignoring single populations that are not consistent with independent clusters). This analysis increased regional differentiation (73%) compared to that of the original regional classification (Table 2). Only 6% of the total variance was found among populations within regions (Table 2). These results were supported by clustering analysis based on UPGMA and DAPC, evidencing a closer evolutionary relationship between Western Colombia and Central American regions than to samples from the Amazonian and northeastern Brazil regions (Figure 4). DAPC showed a clearer differentiation between the three clusters represented, with the first two principal components explaining 91% of the variance in allele frequencies (10 PCs retained). PC1 clearly separates cluster 1 (Western Colombia and Central America) from cluster 2 (Amazonian), while PC2 differentiates these from cluster 3 (Northeastern Brazil; Figure 4A). Consequently, divergence time ranged from 0.1 to 0.2 mya between Pacific and Andean regions, and to 1.4 mya for the most divergent northeastern Brazil (BRA_NE) (Figure 4B)

**Figure 4.**
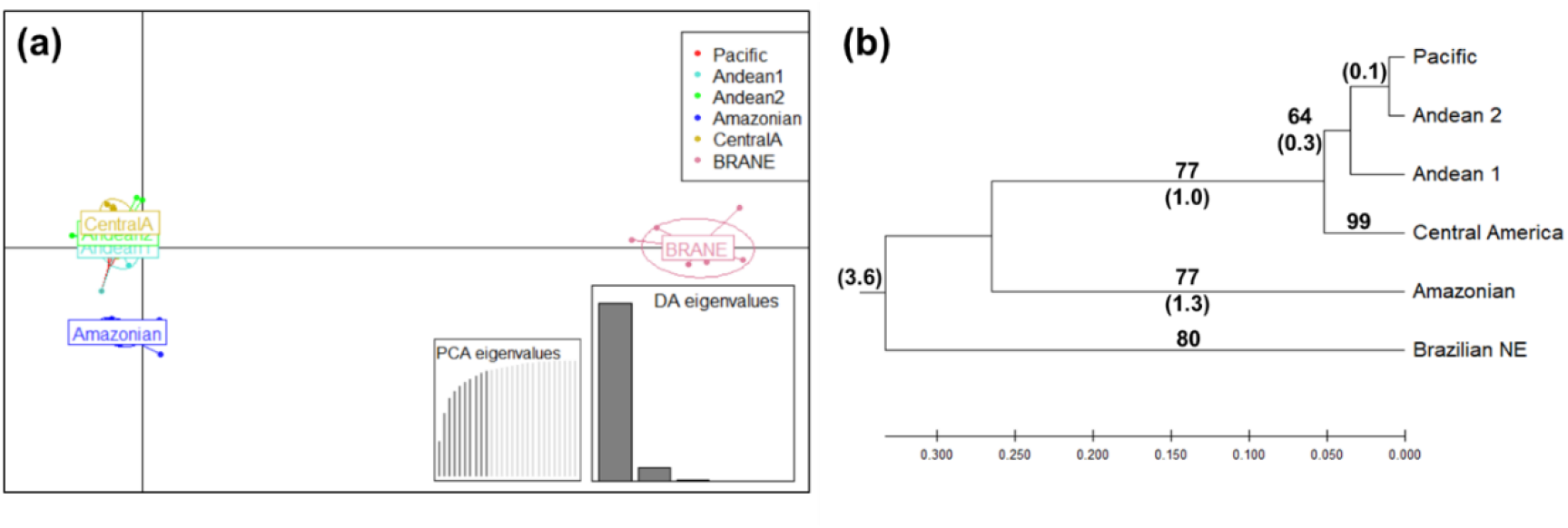
Population genetic structure of the leaf-cutting ant *A. cephalotes* based on clustering analysis. (a) A scatterplot of the discriminant analysis of principal components (DAPC) showing the genetic relationship between of *A. cephalotes* samples. The key describes the colors attributed to each region, and inertia ellipses describe the general distribution of points.Eigenvalues for each PC axis are shown (PC1, vertical; PC2, horizontal). The number of PCA axes retained in each DAPC analysis is shown in the bottom-right inset (gray bars). (b) UPGMA clustering of *mtCOI* sequences from different *A. cephalotes* populations. Support for the branches is given by the bootstrap’s values. Values in parentheses represent divergence time (Mya) between clustered regions.

### 3.3 Scenarios of genetic differentiation – IBD vs IBB

We next used a db-RDA approach combined with model selection to investigate the role played by the Colombian Andes on the regional differentiation detected among major clusters of *A. cephalotes* populations (Western Colombia – WC, Central America – CA and the Amazonian-South America – AS). Two different scenarios were tested by assessing the combination of barriers (the three cordilleras) responsible for isolation. All models were tested by including and excluding northeastern Brazil populations given the particularly high distance of this region compared to the rest of the distribution. The best model following AIC criteria classified populations into two groups in a South-North manner, following the Eastern Colombian range Cordillera partitioning (Figure 5). With this approach, we assessed the relative contribution of IBD and IBB to explain genetic differentiation. Significant IBD was detected following PCNM analysis (Figure 5A).

**Figure 5.**
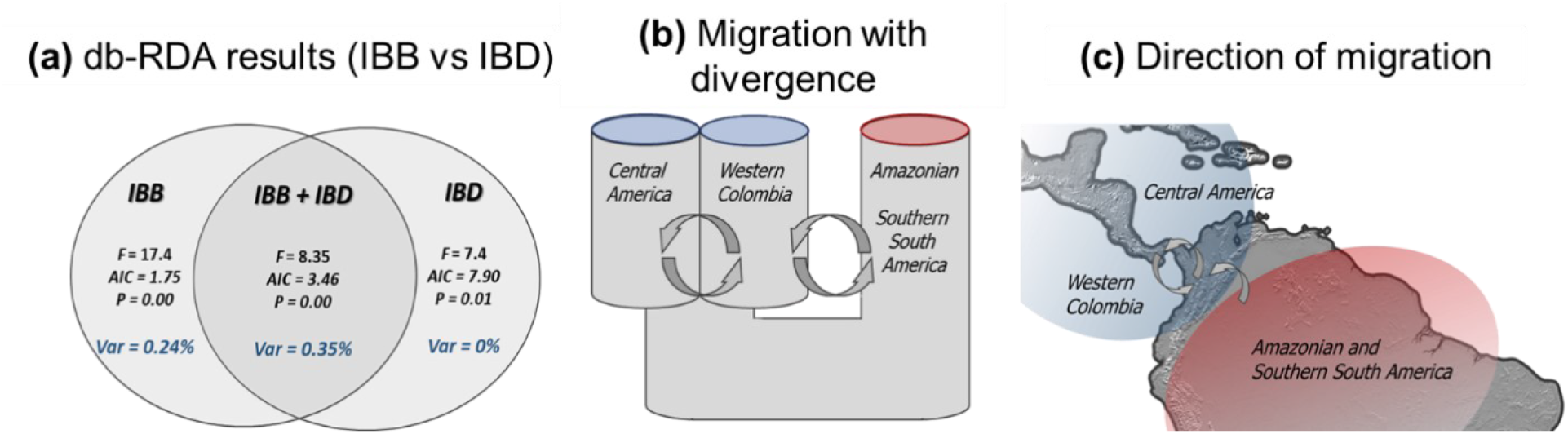
Model comparison of isolating (db-RDA) and migration (Migrate-n) mechanisms driving genetic *mtCOI* differentiation (*Φ*_ST_) in *A. cephalotes*. (a) Variation partitioning analysis based on RDA results of the full model (IBB + IBD) into spatial (IBD: geographical distance) and barrier (IBB: Andes) components. Each circle represents the variation explained for each mechanism, and their overlap the fraction of shared variation. The IBB model remained significant after controlling for IBD (Conditioned model: *F*: 6.44, *P*: 0.003, AIC: 3.46). (b) The divergence (with migration) among the three main genetic groups detected across the Andes [Central America and Western Colombia (blue) and Amazonian-South America (red)]. Model probability = 0.99 among 9 models. (c) The direction of migration model as evaluated in Migrate-n between Central America and Western Colombia (blue) and Amazonian-South America (red). Model probability = 0.99 among 16 models. The rows indicate direction of migration.

When only IBB was considered, a significant contribution of the Andes mountain ranges in shaping patterns of gene flow was observed (Figure 5A), and it remains significant after accounting for IBD. Higher variation explained by the model was detected when both isolating mechanisms (IBD and IBB) were considered, accumulating up to 59% of variation (Figure 5A). This result suggests a combined effect of both the geographical barrier imposed by the Andes as well as geographic distance in shaping genetic differentiation in *A. cephalotes*. Variation partitioning analysis indicated that the best final model showed a strong interaction effect between IBD and IBB that explained 35% of the genetic variation (Figure 5A). The 24% of the variation was partitioned for IBB, and no variation was partitioned for IBD only (Figure 5A).

### 3.4 Migration modelling

Once the genetic structure of *A. cephalotes* populations was confirmed, we used MIGRATE software to investigate the migration pattern that better explain the obtained genetic differentiation across the Andes. As expected from results of population structure, the most likely population model under full Isolation-with-migration confirmed a closer relationship with full migration between Western Colombia – WC and Central America – CA when compared to Amazonian South America – AS. Based on these results, we then fixed a full migration model between WC and CA, and investigated their connection with AS populations by modelling all possible combinations of migration between AS and WC-CA (Figure 5C). The most likely migration model explaining the genetic differentiation of *A. cephalotes* involved unidirectional migration from AS to CA-WC (Posterior probability = 1.0, Figure 5C). Given the historical component of migration models investigated and that AS was found as a more basal group, these two sets of simulations point to southern South America as a more likely origin for *A. cephalotes*.

## 4. DISCUSSION

We took advantage of the geographical distribution of *A. cephalotes* populations in northern Andes (Western Colombia – WC) to investigate their population structure and genetic relationship with ants from Central America – CA and Amazonian –South America – AS. We used a hierarchical approach to investigate the role of the three parallel Andean ranges in northern Andes in shaping this regional differentiation. Here we provide evidence from genetic differentiation and migration modelling demonstrating the role of the Eastern cordillera as an asymmetrical barrier to gene flow for the leaf-cutting ant.

Multiple approaches including Median-Joining network, spatial clustering generated by SAMOVA and pairwise *Φ* _ST_, consistently suggested a North-South genetic differentiation pattern, with the Andes mountains separating two main clades in *A. cephalotes*: 1) WC and CA, 2) Amazonian, as well as a third clade clustering 3) northeastern Brazilian haplotypes. This is consistent with results reported by Solomon et al. (2008), which included a few Colombian samples in similar analyses. Together, these results point to a closer relationship between CA and WC populations and suggest an ancient divergence between these regions and the rest of South America.

The Andean uplift, and particularly the Eastern range define a discreet separation between northern populations (WC and CA) and southern ones (AS), supporting their major role as geographical barriers for *A. cephalotes’* dispersal. Despite the narrowed transition between these two regions, being connected only through the Panama land bridge (Winston, Kronauer, & Moreau, 2016), historical gene flow by female dispersal seems to homogenize genetic variation between them. Female dispersion distance in *A. cephalotes* has been reported being up to 9.7 km (Cherrett, 1968; Helms, 2018), which is a relatively long distance for small insects. Since populations in the Andean region are all found between two of the Colombian ranges (Western and Central), substantial gene flow detected across such geographical barrier may be explained by natural dispersal in between low elevation peaks (< 2,000 m.a.s.l.) (Hernández-Camacho, 1992; Gustavo H. Kattan et al., 2004) in the Western range. However, transport of *A. cephalotes* females mediated by human activities may be a more likely explanation for this dispersion (Montoya-Lerma et al., 2012). Around 70% of human Colombian population transit the Andes, putting intense pressure on natural ecosystems (Cavelier, 1997; G.H. Kattan & Alvarez-Lopez, 1996). High amounts of human traffic across populated areas can facilitate dispersal of *A. cephalotes* individuals across the mountains, allowing admixture with Central American populations when compared with Amazonian populations.

Despite the strong evidence for the Andes mountains as driver of regional differentiation in *A. cephalotes’* populations, we also found evidence for isolation by distance, which makes the interpretation of underlying mechanisms of isolation more difficult to interpret (Crispo, Bentzen, Reznick, Kinnison, & Hendry, 2006; Edwards, Keogh, & Knowles, 2012; Wang et al., 2013). We further investigated both alternative mechanisms of genetic isolation through a db-RDA and model selection approaches (De Queiroz et al., 2017; James et al., 2011; McGaughran, Morgan, & Sommer, 2014; Meirmans, 2015), indicating that both isolating mechanisms have significantly restricted gene flow in *A. cephalotes* populations. The IBB model remains significant after accounting for IBD, which strongly suggest a substantial role of this isolating mechanism in *A. cephalotes*. However, a combined multifactorial effect of both mechanisms was observed, which seems to be a common pattern for heterogenic landscapes (De Queiroz et al., 2017; Guarnizo & Cannatella, 2014; McGaughran et al., 2014; Noguerales et al., 2016), where multiple factors can affect different sections of the species’ range, making it difficult to disentangle their contributions (Crispo et al., 2006; Wang et al., 2013). The Andes ranges in Colombia seem to drive a North-South genetic differentiation in *A. cephalotes*, but geographical distance still has a similar impact within regions on both sides of these mountains.

Most studies on the origin of leaf-cutting ants support their South American origin (Fowler, 1983; Kusnezov, 1963; Mayhé-Nunes & Jaffé, 1998), and so far this continues to be the most accepted hypothesis. This is mainly because the greatest concentration of the leaf-cutting species (Della Lucia et al., 2014; Fernández et al., 2015; Fernández & Sendoya, 2004) and basal Attini species occur in open land biomes of South America (Fowler, 1983; Mayhé-Nunes & Jaffé, 1998). The Central American origin (Branstetter et al., 2017) is mainly based on conditions needed for the evolution of ant agriculture, although such conditions are also possible in southern South America (Kusnezov, 1963; Mueller et al., 2017). Genetic diversity estimates and high divergence of ant populations from the Amazonian and northeastern Brazil regions reflect patterns consistent with ancient populations as expected from a South American origin. This inter-regional diversity is lower than that reported by Lovato (2006) between the *A. cephalotes* populations in the Amazonian and northeastern Brazil. Although both estimations were performed based on standard substitution rates reported for *mtCOI* gene (Hebert et al., 2003), and therefore suffer from the limitations of molecular clock estimates (Hillis, Mable, & Moritz, 1996), this result suggests that *A. cephalotes* populations in South America diverged even before dispersion to Central America. These estimations were consistent with those by Solomon et al. (2008), which places the Amazonian divergence somewhere between the lower Pliocene (4.9 mya) to the mid-Pleistocene (819 kya), with northeastern Brazil populations being more differentiated than Amazonian.

The Amazon basin has been considered a crucial and most recent refugia for many ant species during the Pleistocene glaciation (Bush, 1994; Haffer, 1969; Haffer & Prance, 2001). The Pleistocene refugia hypothesis suggests that historical climate changes occurring during this time, restricted ants’ distribution to the Amazon rainforest (Bush, 1994; Haffer & Prance, 2001). Our results support Solomon et al.(Solomon et al., 2008) findings that non-Amazonian *A. cephalotes* populations diverged during the Pleistocene. This hypothesis predicts multiple Pleistocene refugia, which has previously been proven by paleoclimate analysis (Solomon et al., 2008) and would allow, for example, Pleistocene ant refugia in the mountainous mosaic of Western Colombia.

Together these results strongly support the origin of *A. cephalotes* ants in southern South America, against the Central America alternative. Our simulations of migration models suggest that WC-CA populations in northern South America could only be originated from ancient southern populations. This reveals that the Andean uplift has being an important factor shaping not only the level but also the direction of migration, acting as an asymmetrical barrier to gene flow. Since most evidence point to a South American origin of *A. cephalotes*, these ants managed to across the three Andes mountains ranges from South to Central America. Our results suggest that populations on the west side of the Eastern cordillera come from a secondary introduction from Central America. It is not clear how or when this dispersion occurred, but this cordillera seems to restrict gene flow back to the south in *A. cephalotes*.

Genetic structure in a pest species such as *A. cephalotes* is also affected by several factors other than geographic distance and geographic barriers. Historical and contemporary demographic events and chemical control (Men, Xue, Mu, Hu, & Huang, 2017; Rocha-Olivares & Sandoval-Castillo, 2003; Wei et al., 2013) may also influence population dynamics (Rocha-Olivares & Sandoval-Castillo, 2003). Western Colombian populations exhibit a more heterogenic population structure than others, with low sequence divergence, suggesting ongoing founder effects in this region (Scarpassa, Geurgas, Azeredo-Espin, & Tadei, 2000). Constant pest control in these areas can potentially cause recurrent population extinctions and colonization that are facilitated by human traffic. This might increase population differentiation, while sequence divergence remains low, leading the fixation of unique *mtCOI* lineages (Martins, 2006; Roderick, 1996; Slatkin, 1987).

Our results highlight the importance of studying the transitional lands across the northern Andes when elucidating the role of these mountains in conspecific patterns of historical gene flown in *A. cephalotes*. We detected substantial regional differentiation and provided strong evidence compatible with a South American origin of these ants with subsequent migration to Central America and Western Colombia. The Eastern cordillera seemed to act an asymmetrical barrier to gene flow after this initial dispersion, restricting migration back to southern South America. The genetic dynamics here described also provide insights for pest management strategies since genetic background may reflect differential responses to control plans.

## ACKNOWLEDGEMENTS

We thank Sandra Milena Valencia-Giraldo and Andrea López-Peña for helping with sampling and DNA extraction. Special thanks to Dr. David Johnston-Monje for proofreading the manuscript and editing its scientific English. This work was funded by Vicerrector°a de Investigaciones, Universidad del Valle, Cali, Colombia (grant number: CI71067); and COLCIENCIAS National Program of PhD (grant number: 617-2013).

## AUTHOR CONTRIBUTIONS

VMV participated in the conceptualization, performed ant sampling, molecular methods, data analysis, original draft, reviewed and edited the manuscript. FD participated in data analysis. FD and JML participated in the conceptualization, reviewing and editing of the manuscript. GAVM

performed ant sampling and molecular methods. All authors read and approved the final version of the manuscript.

## REFERENCES

Antonelli, A., Quijada-Mascareñas, A., Crawford, A. J., Bates, J. M., Velazco, P. M., & Wüster, W. (2009). Molecular studies and phylogeography of Amazonian tetrapods and their relation to geological and climatic models. In C. Hoorn & F. Wesselingh (Eds.), Amazonia, landscape and species evolution: A look into the past (pp. 386–404). London, UK: Blackwell Publishing Ltd.

Bandelt, H.-J., Forster, P., & Rohl, A. (1999). Median-joining networks for inferring intraspecific phylogenies. Molecular Biology, 37–48. doi:10.1093/oxfordjournals.molbev.a026036

Beerli, P., Mashayekhi, S., Sadeghi, M., Khodaei, M., & Shaw, K. (2019). Population Genetic Inference With MIGRATE. Current Protocols in Bioinformatics, 68(1), 1–28. doi:10.1002/cpbi.87

Bradburd, G. S., Ralph, P. L., & Coop, G. M. (2013). Disentangling the effects of geographic and ecological isolation on genetic differentiation. Evolution, 67(11), 1–25. doi:10.1111/evo.12193.Disentangling

Branstetter, M. G., Jesovnik, A., Sosa-Calvo, J., Lloyd, M. W., Faircloth, B. C., Brady, S. G., & Schultz, T. R. (2017). Dry habitats were crucibles of domestication in the evolution of agriculture in ants. Proceedings of the Royal Society B, 284, 20170095.

Brumfield, R. T., & Capparella, A. P. (1996). Historical diversification of birds in northwestern South America?: A molecular perspective on the role of vicariant events. Evolution, 50(4), 1607–1624.

Bush, M. B. (1994). Amazonian speciation: A necessarily complex model. Journal of Biogeography, 21(1), 5. doi:10.2307/2845600

Cadena, C. D., Pedraza, C. A., & Brumfield, R. T. (2016). Climate, habitat associations and the potential distributions of Neotropical birds: Implications for diversification across the Andes. Revista de La Academia Colombiana de Ciencias Exactas, Físicas y Naturales, 40(155), 275. doi:10.18257/raccefyn.280

Cavelier, J. (1997). Selvas y bosques montanos. In M.E. Chaves & N. Arango (Eds.), Informe nacional sobre el estado de la biodiversidad. Tomo I (pp. 38–55). Bogotá: Instituto de Investigación de Recursos Biológicos. A. Von Humboldt.

Chacón de Ulloa, P. (1994). Biología e impacto económico de las hormigas. Palmas, 15(4), 25–30.

Chapela, I. H., Rehner, S. ., Schultz, T. R., & Mueller, U. G. (1994). Evolutionary history of the symbiosis between fungus-growing ants and their fungi. Science, 266, 1691–1694.

Chapman, F. M. (1917). The distribution of bird-life in Colombia. Bulletin of American Museum of Natural History, 36, 1–729.

Cherrett, J. (1968). A flight record for queens of Atta cephalotes L. (Hym. Formicidae). Entomologist ‘s Monthly Magazine, 104, 255–256.

Crispo, E., Bentzen, P., Reznick, D. N., Kinnison, M. T., & Hendry, A. P. (2006). The relative influence of natural selection and geography on gene flow in guppies. Molecular Ecology, 15(1), 49–62. doi:10.1111/j.1365-294X.2005.02764.x

De-Silva, D. L., Mota, L. L., Chazot, N., Mallarino, R., Silva-Brandão, K. L., Gómez-Piñerez, L. M., … Elias, M. (2017). North Andean origin and diversification of the largest ithomiine butterfly genus. Scientific Reports, 7, 1–17. doi:10.1038/srep45966

De Queiroz, L. J., Torrente-Vilara, G., Quilodran, C., da Costa Doria, C.R., & Montoya-Burgos, J. I. (2017). Multifactorial genetic divergence processes drive the onset of speciation in an Amazonian fish. PLoS ONE, 12(12), 1–27. doi:10.1371/journal.pone.0189349

Della Lucia, T. M., Gandra, L. C., & Guedes, R. N. (2014). Managing leaf-cutting ants: Peculiarities, trends and challenges. Pest Management Science, 70(1), 14–23. doi:10.1002/ps.3660

Dick, C. W., Roubik, D. W., Gruber, K. F., & Bermingham, E. (2004). Long-distance gene flow and cross-Andean dispersal of lowland rainforest bees (Apidae: Euglossini) revealed by comparative mitochondrial DNA phylogeography. Molecular Ecology, 13(12), 3775–3785. doi:10.1111/j.1365-294X.2004.02374.x

Dupanloup, I., Schneider, S., & Excoffier, L. (2002). A simulated annealing approach to define the genetic structure of populations. Molecular Ecology, 11, 2571–2581.

Edwards, D. L., Keogh, J. S., & Knowles, L. L. (2012). Effects of vicariant barriers, habitat stability, population isolation and environmental features on species divergence in the south-western Australian coastal reptile community. Molecular Ecology, 21(15), 3809–3822. doi:10.1111/j.1365-294X.2012.05637.x

Elias, M., Joron, M., Willmott, K., Silva-BrandÃo, K. L., Kaiser, V., Arias, C. F., … Jiggins, C. D. (2009). Out of the Andes: Patterns of diversification in clearwing butterflies. Molecular Ecology, 18(8), 1716–1729. doi:10.1111/j.1365-294X.2009.04149.x

Excoffier, L., & Lischer, H. E. L. (2010). Arlequin suite ver 3.5: A new series of programs to perform population genetics analyses under Linux and Windows. Molecular Ecology Resources, 10(3), 564–567. doi:10.1111/j.1755-0998.2010.02847.x

Fernández, F., Castro-Huertas, V., & Serna, F. (2015). Hormigas cortadoras de hojas de Colombia: Acromyrmex & Atta (Hymenoptera: Formicidae). Fauna de Colombia (Monografía). Bogotá: Instituto de Ciencias Naturales, Universidad Nacional de Colombia.

Fernández, F., & Sendoya, S. (2004). List of Neotropical ants (Hymenoptera: Formicidae). Revista Biota Colombiana, 5(1), 3–93. doi:10.1017/CBO9781107415324.004

Fowler, H. (1983). Latitudinal gradients and diversity of the leaf-cutting ants (Atta and Acromyrmex) (Hymenoptera: Formicidae). Revista de Biología Tropical, 31(2), 213–216.

Fu, Y.-X. (1997). Statistical test of neutrality of mutations against population growth, hitchhiking and background selection. Genetics, 147, 915–925.

Guarnizo, C. E., & Cannatella, D. C. (2014). Geographic determinants of gene flow in two sister species of tropical andean frogs. Journal of Heredity, 105(2), 216–225. doi:10.1093/jhered/est092

Haffer, J. (1969). Speciation in Amazanian forest birds. Science, 165(3889), 131–137.

Haffer, J., & Prance, G. T. (2001). Climatic forcing of evolution in Amazonia during the Cenozoic: On the refuge theory of biotic differentiation. Amazoniana, 16, 579–607. doi:10.1371/journal.pntd.0000502

Harrison, R. G. (1991). Molecular Changes at Speciation. Annual Review of Ecology and Systematics, 22, 281–308.

Hebert, P. D. N., Cywinska, A., Ball, S. L., & Waard, J. R. (2003). Biological identifications through DNA barcodes. Ingenieria e Investigacion, 270, 313–321. doi:10.1098/rspb.2002.2218

Helms, J. (2018). The flight ecology of ants (Hymenoptera: Formicidae). Myrmecological News, 26, 19–30. doi:10.25849/myrmecol.news

Hernández-Camacho, J. (1992). Caracterización geográfica de Colombia. In G. Halffter (Ed.), La diversidad biológica de Iberoamérica I (pp. 45–52). Acta Zoológica Mexicana.

Hillis, D. M., Mable, B., & Moritz, C. (1996). Applications of molecular systematics: the state of the field on a look to the future. In: D. M. Hillis, C. Moritz, & B. Mable (Eds.), Molecular systematics (2nd ed., pp. 515–543). Sinauer, Sunderland.

Hoorn, C., Wesselingh, F. P., Ter Steege, H., Bermudez, M. A., Mora, A., Sevink, J., … Antonelli, A. (2010). Amazonia through time: Andean uplift, climate change, landscape evolution, and biodiversity. Science, 330, 927–931. doi:10.1126/science.1194585

James, P. M. A., Coltman, D. W., Murray, B. W., Hamelin, R. C., & Sperling, F. A. H. (2011). Spatial genetic structure of a symbiotic beetle-fungal system: Toward multi-taxa integrated landscape genetics. PLoS ONE, 6(10), 1–11. doi:10.1371/journal.pone.0025359

Jombart, T. (2008). Adegenet: A R package for the multivariate analysis of genetic markers. Bioinformatics, 24(11), 1403–1405. doi:10.1093/bioinformatics/btn129

Jombart, T., Devillard, S., & Balloux, F. (2010). Discriminant analysis of principal components: a new method for the analysis of genetically structured populations. BMC Genetics, 11(94), 1–15. doi:10.1371/journal.pcbi.1000455

Kattan, G. H., & Alvarez-Lopez, H. (1996). Preservation and management of biodiversity in fragmented landscapes in the Colombian Andes. In J. Schelhas & R. Greenberg (Eds.), Forest patches in tropical landscapes (pp. 3–18). Washington D.C. Island Press.

Kattan, Gustavo H., Franco, P., Rojas, V., & Morales, G. (2004). Biological diversification in a complex region: A spatial analysis of faunistic diversity and biogeography of the Andes of Colombia. Journal of Biogeography, 31(11), 1829–1839. doi:10.1111/j.1365-2699.2004.01109.x

Kronauer, D. J. C., Hölldobler, B., & Gadau, J. (2004). Phylogenetics of the new world honey ants (genus Myrmecocystus) estimated from mitochondrial DNA sequences. Molecular Phylogenetics and Evolution, 32(1), 416–421. doi:10.1016/j.ympev.2004.03.011

Kusnezov, N. (1963). Zoogeografía de las hormigas en Suramérica. Acta Zoológica Lilloana, 19, 25–186.

Lagomarsino, L. P., Condamine, F. L., Antonelli, A., Mulch, A., & Davis, C. C. (2016). The abiotic and biotic drivers of rapid diversification in Andean bellflowers (Campanulaceae). New Phytologist, 210, 1430–1442. doi:10.1111/nph.13920

Leal, I. R., Wirth, R., & Tabarelli, M. (2014). The multiple impacts of leaf-cutting ants and their novel ecological role in human-modified neotropical forests. Biotropica, 46(5), 516–528. doi:10.1111/btp.12126

Legendre, P., & Fortin, M. J. (2010). Comparison of the Mantel test and alternative approaches for detecting complex multivariate relationships in the spatial analysis of genetic data. Molecular Ecology Resources, 10(5), 831–844. doi:10.1111/j.1755-0998.2010.02866.x

Librado, P., & Rozas, J. (2009). DnaSP v5: A software for comprehensive analysis of DNA polymorphism data. Bioinformatics, 25(11), 1451–1452. doi:10.1093/bioinformatics/btp187

Lovato, L. (2006). Estudos morfologicos e analises de seqüências do gene mitocondrial Citocromo Oxidase I (COI) em populações de Atta cephalotes (L., 1758) (Doctoral dissertation). Retrieved from Biblioteca Universidade do Vale do Paraíba.

Luebert, F., & Weigend, M. (2014). Phylogenetic insights into Andean plant diversification. Frontiers in Ecology and Evolution, 2(27), 1–17. doi:10.3389/fevo.2014.00027

Martins, J. J. (2006). Evolução de genes e pseudogenes mitocondriais em Atta cephalotes (Linnaeus, 1758) (Master’s thesis). Retrieved from repositorio of Universidad Estatal Paulista, Unesp.

Masello, J. F., Quillfeldt, P., Munimanda, G. K., Klauke, N., Segelbacher, G., Schaefer, H. M., … Moodley, Y. (2011). The high Andes, gene flow and a stable hybrid zone shape the genetic structure of a wide-ranging South American parrot. Frontiers in Zoology, 8(June). doi:10.1186/1742-9994-8-16

Mayhé-Nunes, A. J., & Jaffé, K. (1998). On the biography of Attini (Hymenoptera: Formicidae). Ecotropicos, 11(1), 45–54.

McGaughran, A., Morgan, K., & Sommer, R. J. (2014). Environmental variables explain genetic structure in a beetle-associated nematode. PLoS ONE, 9(1), 1–15. doi:10.1371/journal.pone.0087317

Meirmans, P. G. (2015). Seven common mistakes in population genetics and how to avoid them. Molecular Ecology, 24, 3223–3231.

Men, Q., Xue, G., Mu, D., Hu, Q., & Huang, M. (2017). Mitochondrial DNA markers reveal high genetic diversity and strong genetic differentiation in populations of Dendrolimus kikuchii Matsumura (Lepidoptera: Lasiocampidae). PLoS ONE, 12(6), 1–16. doi:10.1371/journal.pone.0179706

Milá, B., Wayne, R. K., Fitze, P., & Smith, T. B. (2009). Divergence with gene flow and fine-scale phylogeographical structure in the wedge-billed woodcreeper, Glyphorynchus spirurus, a neotropical rainforest bird. Molecular Ecology, 18(14), 2979–2995. doi:10.1111/j.1365-294X.2009.04251.x

Miller, M. J., Bermingham, E., Klicka, J., Escalante, P., Amaral, F. S. R., Weir, J. T., & Winker, K. (2008). Out of Amazonia again and again: episodic crossing of the Andes promotes diversification in a lowland forest flycatcher. Proceedings of the Royal Society of London B, 275, 1133–1142.

Montoya-Lerma, J., Giraldo-Echeverri, C., Armbrecht, I., Farji-Brener, A., & Calle, Z. (2012). Leaf-cutting ants revisited: Towards rational management and control. International Journal of Pest Management, 58(3), 225–247. doi:10.1080/09670874.2012.663946

Mueller, U. G., Ishak, H. D., Bruschi, S. M., Smith, C. C., Herman, J. J., Solomon, S. E., … Bacci, M.J. (2017). Biogeography of mutualistic fungi cultivated by leafcutter ants. Molecular Ecology, 26(24), 6921–6937.

Noguerales, V., Cordero, P. J., & Ortego, J. (2016). Hierarchical genetic structure shaped by topography in a narrow-endemic montane grasshopper. BMC Evolutionary Biology, 16(96), 1–15. doi:10.1186/s12862-016-0663-7

Oksanen, J., Blanchet, F. G., Friendly, M., Kindt, R., Legendre, P., McGlinn, D., … Wagner, H. (2019). Vegan: Community Ecology Package. R Package Version 2.2-0. Community ecology package. Retrieved from https://cran.r-project.org/web/packages/vegan/vegan.pdf

Pérez-Escobar, O. A., Gottschling, M., Chomicki, G., Condamine, F. L., Klitgård, B. B., Pansarin, E., & Gerlach, G. (2017). Andean mountain building did not preclude dispersal of lowland epiphytic orchids in the Neotropics. Scientific Reports, 7(1), 1–10. doi:10.1038/s41598-017-04261-z

Ramos-Onsins, S. E., & Rozas, J. (2002). Statistical properties of new neutrality tests against population growth. Molecular Biology and Evolution, 19(12), 2092–2100. doi:Doi 10.1093/Molbev/Msl052

Rocha-Olivares, A., & Sandoval-Castillo, J. R. (2003). Mitochondrial diversity and genetic structure in allopatric populations of the Pacific red snapper Lutjanus peru. Ciencias Marinas, 29(2), 197–209.

Roderick, G. K. (1996). Geographic structure of insect populations: gene flow, phylogeography, and their uses. Annual Review of Entomology, 41, 325–352.

Salgado-Roa, F. C., Pardo-Diaz, C., Lasso, E., Arias, C. F., Solferini, V. N., & Salazar, C. (2018). Gene flow and Andean uplift shape the diversification of Gasteracantha cancriformis (Araneae: Araneidae) in Northern South America. Ecology and Evolution, 8(14), 7131–7142. doi:10.1002/ece3.4237

Scarpassa, V. M., Geurgas, S., Azeredo-Espin, A. M. L., & Tadei, W. P. (2000). Genetic divergence in mitochondrial DNA of Anopheles nuneztovari (Diptera: Culicidae) from Brazil and Colombia. Genetics and Molecular Biology, 23(1), 71–78. doi:10.1590/S1415-47572000000100013

Shi, W., Kerdelhué, C., & Ye, H. (2012). Genetic structure and inferences on potential source areas for Bactrocera dorsalis (Hendel) based on mitochondrial and microsatellite markers. PLoS ONE, 7(5), 1–15. doi:10.1371/journal.pone.0037083

Simon, C., Frati, F., Beckenbach, A., Crespi, B., Liu, H., & Flook, P. (1994). Evolution, weighting, and phylogenetic utility of mitochondrial gene sequences and a compilation of conserved polymerase chain reaction primers. Annals of the Entomological Society of America, 87(6), 651–701. doi:10.1093/aesa/87.6.651

Slatkin, M. (1987). Gene flow and the geographic structure of natural populations. Science, 236(4803), 787–792. Retrieved from https://www.jstor.org/stable/1699930

Solomon, S. E., Bacci, M., Martins, J., Vinha, G. G., & Mueller, U. G. (2008). Paleodistributions and comparative molecular phylogeography of leafcutter ants (Atta spp.) provide new insight into the origins of Amazonian diversity. PLoS ONE, 3(7). doi:10.1371/journal.pone.0002738

Tajima, F. (1989). Statistical method for testing the neutral mutation hypothesis by DNA polymorphism. Genetics, 123, 585–595.

Tamura, K., Stecher, G., Peterson, D., Filipski, A., & Kumar, S. (2013). MEGA6: Molecular evolutionary genetics analysis version 6.0. Molecular Biology and Evolution, 30(12), 2725– 2729. doi:10.1093/molbev/mst197

Wang, I. J., Glor, R. E., & Losos, J. B. (2013). Quantifying the roles of ecology and geography in spatial genetic divergence. Ecology Letters, 16(2), 175–182. doi:10.1111/ele.12025

Weber, N. A. (1972). Gardening ants, the attines. Memoirs of the American Philosophical Society (Vol. 92).

Wei, S. J., Shi, B. C., Gong, Y. J., Jin, G. H., Chen, X. X., & Meng, X. F. (2013). Genetic structure and demographic history reveal migration of the diamondback moth Plutella xylostella (Lepidoptera: Plutellidae) from the southern to northern regions of China. PLoS ONE, 8(4), 1–14. doi:10.1371/journal.pone.0059654

Wetterer, J. K., Schultz, T. R., & Meier, R. (1998). Phylogeny of fungus-growing ants (Tribe Attini) based on mtDNA sequence and morphology. Molecular Phylogenetics and Evolution, 9(1), 42–47. doi:10.1006/mpev.1997.0466

Winston, M. E., Kronauer, D. J. C., & Moreau, C. S. (2016). Early and dynamic colonization of Central America drives speciation in Neotropical army ants. Molecular Ecology, 26(3), 859– 870. doi:10.1111/mec.13846

